# Using Genetic Information to Improve the Prediction of Individual Food Choice: A Case Study of Alcoholic Beverages

**DOI:** 10.1101/2020.12.29.424769

**Authors:** Chen Zhu, Timothy Beatty, Qiran Zhao, Wei Si, Qihui Chen

**Affiliations:** College of Economics and Management, China Agricultural University, Qinghuadonglu No. 17, Beijing, 100083, China; China Center for Genoeconomic Studies (CCGS), Beijing, 100083, China; Department of Agricultural and Resource Economics, UC Davis, CA 95616, USA

**Keywords:** Consumer preference, food choice, choice experiment, genetic factor, machine learning

## Abstract

Individual food choices and consumption are closely relating to one’s diet, nutrition, and health. Using the case of alcoholic beverages, this study extends the random-utility framework by incorporating genetic information into consumer demand models, and demonstrates the significant impact of genetic factors on individual food choice decisions in a novel way. Integrating individual-level responses of discrete choice experiments (DCE), genotyping data, and socioeconomic/demographic characteristics of 484 participants collected from face-to-face interviews in mainland China, we employ a machine learning-based classification (MLC) approach to identify and predict individual choices. We show that genetic factors are critical to explaining variations in both general drinking behavior and choices of particular products. We systematically compared the performance of traditional discrete choice models and MLC models without and with genetic factors. The MLC predictive model with both socio-demographic and genetic features yields the highest accuracy of 74.7% and AUC-ROC of 0.85. Our findings warrant further economic studies of human behaviors with the integration of genetic data.

## 1. Introduction

Food choice and intake have profound effects on an individual’s dietary, nutritional, and health outcomes. Not only is a better understanding of the determinants of individual food choice the foundation of accurate prediction of individual food consumption behavior, but it may also facilitate the present developments of precision nutrition and precision marketing (Arora et al., 2008; Garcia-Perez et al., 2020). A large body of evidence from economics, marketing, and consumer behavior have shown that product attributes (e.g., price and nutrition contents) and individual socioeconomic/demographic characteristics (e.g., age, gender, education, and income) are the most significant predictors of individual choice behavior (Ansari et al., 2000; Shepherd, 1999). However, recent breakthroughs in genetics and nutritional science have identified a number of novel genetic variants that are robustly associated with individual food intake and dietary patterns (Matoba et al., 2020; Cole et al., 2020; Meddens et al., 2020). This highlights the possible role of genetic factors as one of the critical determinants of individual food choice decisions.

Despite the potential importance of genetic determinants of individuals’ food choices, the application of genetic factors in predicting individuals’ decision-making behavior is still scarce. The paucity of individual-level genetic data combined with consumer behavior data and communication barriers across disciplines render this line of research extremely challenging. Earlier tools for predicting consumer behavior, such as those based on discrete choice models and conjoint analyses, are subject to a relatively low predictive power (McFadden, 1986; Fader and Hardie, 1996). Nevertheless, recent developments in machine learning techniques provide promising prospects to greatly enhance the efficiency and accuracy of individual decision-making predictions (West et al., 1997; Alon et al., 2001; Cui and Curry, 2005; Lemmens and Croux, 2006; Haaf et al., 2014; Lusk, 2017; Shao et al., 2020).

In this study, we assess how genetic factors perform in predicting individual choice of food products, in comparison with conventional socioeconomic/demographic attributes. We conducted a discrete choice experiment (DCE) integrating novel genetic information in a state-of-the-art machine learning framework. DCE is an advanced economic research design that can reliably elicit individuals’ true preferences and choice intentions by replicating real-life situations (Lusk et al., 2003; Mangham et al., 2009). This study attempts to answer the following research questions: whether genetic information can help explain individual food choice behavior? If so, to what extent the prediction accuracy of individual choice can be improved by such an integration?

The food product under consideration is the alcoholic beverage, which is appropriately pertinent to our research question for three reasons. First, apart from its unique social and cultural values (Haucap and Herr, 2013), alcohol drinking has been found to be associated with a large number of adverse health conditions and has been extensively investigated in various disciplines (Dietler, 2006; Sudhinaraset et al., 2016; Griswold et al., 2018). Second, the alcoholic beverage market is highly differentiated by product attributes and flavors, which makes this product particularly suitable for investigating consumer heterogeneity arising from conventional socio-demographic and novel genetic variations (Rojas and Peterson, 2008). Finally, the heritability of alcohol consumption behavior, in general, is estimated to be approximately 40% (Vrieze et al., 2014; Clarke et al., 2017), which provides scientific prerequisites for the potential role that genetic factors may play in determining one’s choice of a specific alcoholic product.

By linking three sets of data, i.e., data from the DCE, individual genotyping data, and traditional socio-demographic information of 484 participants collected from face-to-face interviews in mainland China, we employed a machine learning-based classification (MLC) approach to assess how well genetic information can aid in predicting individual food choice decisions. The DCE we designed incorporates three main multiple-level attributes: alcohol type, price, and amount. Building on the existing literature, we train the MLC models with three different sets of individual-level features: socioeconomic/demographic attributes (e.g., age, gender, income), single genetic variants and polygenic scores (PGSs) of complex traits. Figure 1 illustrates a schematic of the study design.

**Fig. 1.**
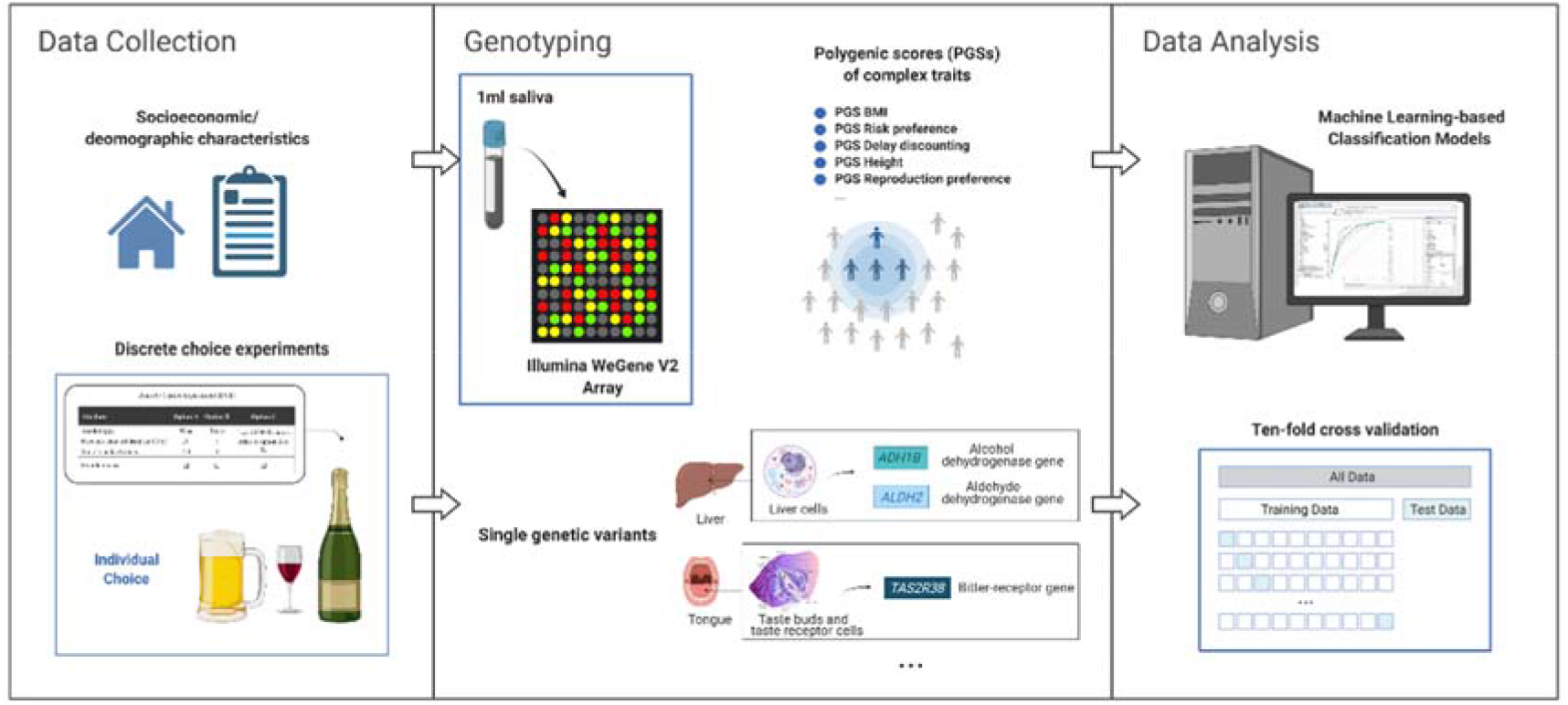
Illustration of the research design. Notes: Key features used in the machine learning-based classification models include the following sets: (1) product attributes; (2) demographic and socioeconomic characteristics; (3) single genetic markers; and (4) polygenic scores of complex human traits. Source: Author’s design.

The remaining of this paper is structured as follows. Section 2 briefly introduces how genetic information can be conceptually incorporated into the random-utility framework of consumer demand models. Section 3 details the implementation of the discrete choice experiment and the proposed machine learning methodology. Section 4 presents and discusses empirical results and implications. Section 5 concludes.

## 2. Incorporating Genetic Variables into Consumer Demand Models

We model consumer choices based on the random-utility theory (McFadden, 1986), and incorporate the individual-level genetic information to further explain consumer heterogeneity besides conventional socio-demographic characteristics. A well-known flexible model used to analyze consumer demand is the mixed logit model.

Let consumer *i* chooses product *j* in choice situation *t* to maximize the conditional indirect utility:

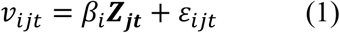

where ***Z_jt_*** contains observed product attributes, such as price and nutritional quality. *β_i_* is a vector of individual-specific parameters. *ε_ijt_* is an independent and identically distributed disturbance term. To capture consumer heterogeneity, we specify the random coefficient *β_i_* to be composed of fixed and variable components that variate across conventional socio-demographic factors ***D_i_***, novel genetic factors ***G_i_***, and unobserved characteristics *ξ_i_* of each consumer:

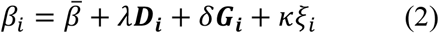

where 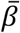 measures the mean preference that is common to all consumers. *λ* and *δ* are matrices with elements measuring how consumers’ preferences vary with conventional socio-demographics and novel genetics, respectively. *ξ_i_* follows a standard multivariate normal distribution with *κ* as a scaling vector. Consequently, the probability that consumer *i* chooses product *j* in choice situation *t* is:

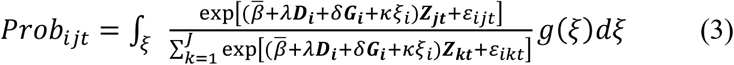

Hence, the random-utility model of consumer demand can be naturally extended to incorporate individual-level genetic information as shown in equation (3). Notice that the genetic factors, ***G_i_***, are determined at conception and remain stable through an individual’s lifetime, and thus can be modeled mostly the same as other demographic variables such as gender and ethnicity. The relative impact of genetic variables may vary by the product category studied.

## 3. Data and Methods

### 3.1 Data collection

The survey was designed and implemented by the Center for Human Capital and Genoeconomics (CHCG) at China Agricultural University in the summer of 2019 (Zhu et al., 2020). The Institutional Review Board of China Agricultural University approved the protocol. Fifty villages from seven provinces in China (Zhejiang, Shandong, Anhui, Henan, Heilongjiang, Yunnan and Xinjiang) were selected; in each village 10 households were then randomly selected. Prior to data collection, all participants signed an informed consent form after receiving a careful explanation about the purpose of this study. All participants were informed that their responses were completely voluntary and confidential, and were invited to contact the research team later if they had any further questions regarding any aspect of the study. Table 1 presents summary statistics of characteristics of the 484 participants of the survey whose responses met the criteria for quality control.

**Table 1.** Summary statistics of the analytic sample (n = 484)

| Variable | Mean/Percentage | S.D. |
| --- | --- | --- |
| *Socioeconomic/demographic characteristics:* |  |  |
| Age | 49.5 | 11.8 |
| Years of schooling | 8.2 | 3.5 |
| Annual household income (in CNY 10,000) | 7.5 | 12.8 |
| Household size | 4.3 | 1.8 |
| No. of books in the house when at age of 8<sup>a</sup> | 8.3 | 23.4 |
| Individualism<sup>b</sup> | 0.29 | 0.26 |
| No. of friends that talk frequently<sup>c</sup> | 6.1 | 3.5 |
| Gender | Male: 73.2%; Female: 26.8% |  |
| Party member | No: 66.4% |  |
| Ethnic minority | Yes: 19.5%; No: 80.5% |  |
| General drinking frequency | Never: 54.0%; Occasionally/Frequently: 46.0% |  |
| Parental drinking frequency | Never: 32.2%; Occasionally/Frequently: 67.8% |  |
| Single genetic variants: |  |  |
| ALDH2 rs671 | AA: 4.1%; AG: 29.7%; GG: 66.2% |  |
| ADH1B rs1229984 | AA: 41.7%; AG: 44.3%; GG: 14.0% |  |
| TAS2R38 rs1726866 | AA: 11.2%; AG: 44.7%; GG: 44.1% |  |
| CYP1A2 rs762551 | AA: 40.7%; AC: 44.5%; CC: 14.8% |  |
| OXTR rs53576 | AA: 52.0%; AG: 38.9%; GG: 9.1% |  |
| LRRTM4 rs11126630 | TT: 25.6%; TC: 47.3%; CC: 27.1% |  |
| Polygenic scores of complex traits: |  |  |
| Height | -1.05 | 0.44 |
| Body mass index (BMI) | 1.01 | 0.35 |
| Educational attainment (EA) | 0.30 | 0.08 |
| Risk preference | 0.03 | 0.07 |
| Reproduction preference | 0.04 | 0.16 |
| Delay discounting | 0.03 | 0.15 |
| Risk of depression | 0.04 | 0.12 |

Source: Author's survey. Note: <sup>a</sup>. The total number of books in the house at the age of eight was used as an indicator of the participant's family background in early childhood and adolescent (Hanushek and Woessmann, 2011). <sup>b</sup>. An individualism score constructed from the triad task was used to measure the participant's cultural thought (Talhelm et al., 2014). <sup>c</sup>. Social relation was measured by the number of friends that the participant frequently communicated with (Zhang and Zhao, 2015).

### 3.2 Genetic factors candidates

We start with a few plausible genetic factors candidates that might be connected to individual food preference, choice, and intake. Since this is an initial exploration, we constrain the set to genetic factors that fit two criteria: (i.) factors that have received multiple empirical verifications that linked to diet, metabolism, taste perception, and substance use (esp. alcohol); (ii.) factors that give a broad coverage of physiological differences and behavioral traits. This resulted in six single genetic variants for alcohol metabolism capacity (*ALDH2* rs671 and *ADH1B* rs1229984), bitter taste perception (*TAS2R38* rs1726866), caffeine metabolism capacity (*CYP1A2* rs762551), and personality (*OXTR* rs53576 and *LRRTM4* rs11126630); as well as seven polygenic scores for bodily dimensions (PGS height and PGS BMI), mental health (PGS risk of depression), economic preferences (PGS risk preference, PGS delay discounting, and PGS reproduction preference), and socioeconomic outcome (educational attainment). Table 2 lays out all genetic variables, related phenotypes, literature sources, and value interpretation.

**Table 2.** Description and Interpretation of Genetic Factors

| Genetic Factor | Related Phenotype | Reference | Variable Type and Value | Value Interpretation |
| --- | --- | --- | --- | --- |
| ***Single genetic variants:*** |  |  |  |  |
| *ALDH2* rs671<br>(Aldehyde dehydrogenase 2 gene) | Alcohol metabolism capacity | Edenberg and McClintick, 2018 | Discrete: 0 (genotype=GG), 1 (AG), 2 (AA) | Higher values linked to reduced metabolism of ethanol and alcohol intolerance/aversive reactions due to aldehyde dehydrogenase deficiency. |
| *ADH1B* rs1229984<br>(Alcohol dehydrogenase 1 B gene) | Alcohol metabolism capacity | Edenberg and McClintick, 2018 | Discrete: 0 (genotype=GG), 1 (AG), 2 (AA) | Higher values linked to reduced metabolism of ethanol and alcohol intolerance/aversive reactions due to higher activity of alcohol dehydrogenase. |
| *TAS2R38* rs1726866<br>(Taste receptor 2 member 38 gene) | Bitter taste | Risso et al., 2016 | Discrete: 0<br>(genotype=TT), 1<br>(CT/CC) | Higher values linked to higher bitter taste sensitivity. |
| *CYP1A2* rs762551<br>(Cytochrome P450 1A2 gene) | Caffeine metabolism capacity | Denden et al., 2016 | Discrete: 0<br>(genotype=CC/AC), 1 (AA) | Higher values linked to increased metabolism of caffeine. |
| *OXTR* rs53576<br>(Oxytocin receptor gene) | Oxytocin | Li et al., 2015 | Discrete: 0<br>(genotype=AA/AG), 1 (GG) | Higher values linked to emotional traits of increased optimistic and empathetic. |
| *LRRTM4* rs11126630<br>(Leucine-rich repeat transmembrane neuronal 4 gene) | Childhood aggressive behaviour | Pappa et al., 2016 | Discrete: 0<br>(genotype=CC), 1 (CT), 2 (TT) | Higher values linked to more aggressive behaviour in childhood. |
| ***Polygenic scores of complex traits:*** |  |  |  |  |
| PGS height | Body height | Wood et al., 2014 | Continuous | Higher values linked to taller height. |
| PGS BMI | Body mass index | Yengo et al., 2018 | Continuous | Higher values linked to higher BMI. |
| PGS EA | Educational attainment | Lee et al., 2018 | Continuous | Higher values linked to higher levels of educational attainment. |
| PGS risk preference | Risk preference | Linnér et al., 2019 | Continuous | Higher values linked to more risk-loving. |
| PGS reproduction preference | Age at first birth | Barban et al., 2016 | Continuous | Higher values linked to later age at first birth (i.e., having baby at an older age). |
| PGS delay discounting | Delay discounting | Sanchez-Roige et al., 2018 | Continuous | Higher values linked to higher delay discounting (i.e., preferring instant gratification). |
| PGS depression | Risk of depression | Wray et al., 2018 | Continuous | Higher values linked to greater risk of depression. |

Source: Author's summary.

### 3.3 Design and implementation of the discrete choice experiment

A discrete choice experiment (DCE) was conducted to evaluate the participants’ preference of alcoholic drinks (Kanninen, 2002). Based on results of a pilot study, we selected three multiple-level attributes in the DCE: alcohol type (alcopop, beer, wine, or spirits), price (CNY 5, 20, or 50 per standard drink), and amount (1, 2-4, or 5 standard drinks). Because the aforementioned attributes yield a total of 36 (= 4×3×3) possible products, a D-efficient fractional factorial experimental design was used to select ten choice scenarios to be used in the actual DCE. As illustrated by the sample choice scenario (Table 3), each participant was presented with the same ten choice sets/scenarios with three options in each scenario, two featuring alcoholic products that have close market substitutes and one no-purchasing (or opt-out) option. The inclusion of the no-purchasing or opt-out option is crucial as it adds additional realism to the choice task and consumer behavior forecasting (Lancsar et al., 2017).

**Table 3.**
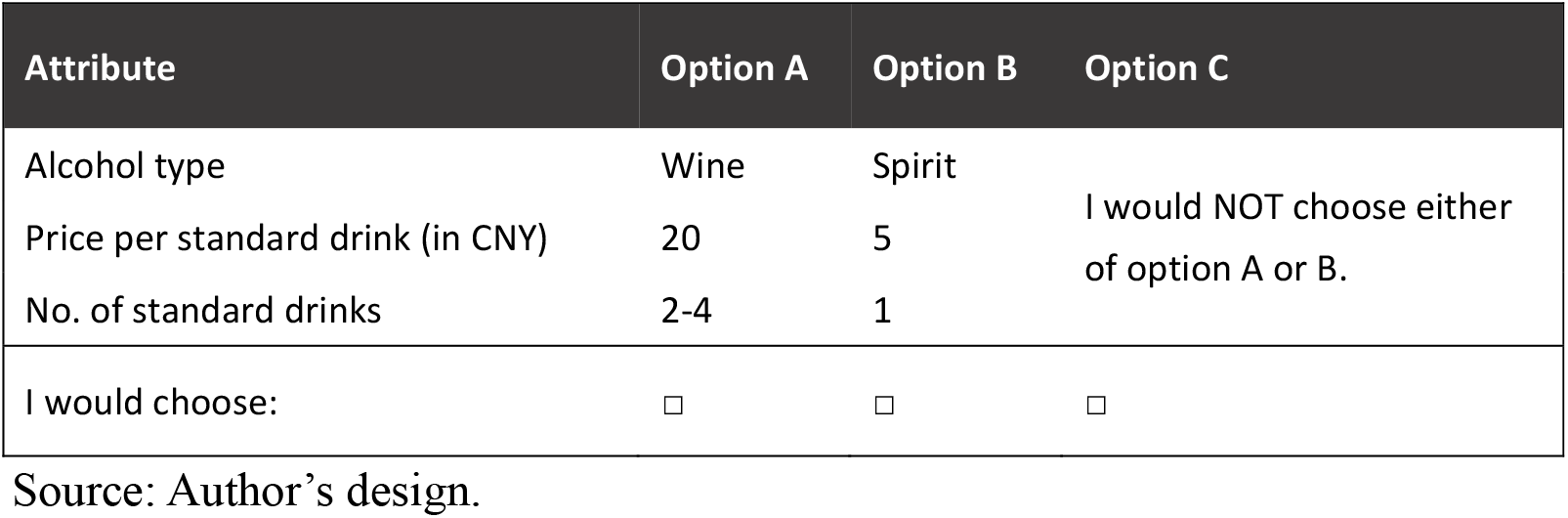
An example of a choice scenario in DCE

### 3.4 Genotyping and quality controls

1 ml saliva samples were collected from all participants during the face-to-face interview. DNA was extracted from these saliva samples using the Illumina WeGene V2 Array. For each individual, we obtained more than 10 million Single Nucleotide Polymorphisms (SNPs), which were then used to construct six single genetic markers (i.e., rs1229984, rs671, rs1726866, rs762551, rs53576, and rs11126630) and seven polygenic scores of genome-wide complex traits (i.e., height, BMI, educational attainment, risk preference, reproduction preference, delay discounting, and risk of depression; see Table 1). Polygenic scores (also known as “polygenic risk scores”) are aggregated effects of hundreds and thousands of trait-associated DNA variants identified in genome-wide association studies (GWAS) and can be used to predict propensities toward certain traits and behavior outcomes at the individual level (Krapohl et al., 2018; Wang et al., 2020). While existing GWAS were mostly based on samples of European ancestry, recent studies indicate that their results can well apply to East Asian (e.g., Chinese) populations (Duncan et al., 2019). Excluding individuals who did not pass the quality control yielded a total of 484 participants with linked phenotyping and genotyping data. This yields a statistical sample of 4,840 observations (484 individuals × 10 choice sets).

### 3.5 Machine learning based-classification models

Given our goal to predict participants’ choice of alcoholic beverage, the outcome variable of the multi-class classification task was defined as follows: Class 1 if the respondent chooses an alcopop product; Class 2 if the respondent chooses a beer product; Class 3 if the respondent chooses a wine product; Class 4 if the respondent chooses a spirit product; and Class 5 if the respondent chooses the opt-out option. The classification task requires a target function *f* that maps an input vector *x* (i.e., features) onto the corresponding predefined class labels of *y*.

The extreme gradient boosting (Chen and Guestrin, 2016), an advanced supervised ensemble approach, was adopted to build and train our machine learning based-classification model. The extreme gradient boosting is built in a stage-wise way and uses a more regularized model formalization to minimize overfitting and improve generalizability. Formally, one can learn the multi-class classification model by minimizing the regularized loss function over a training set containing *n* observations:

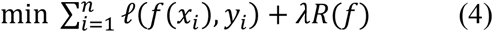

where *ℓ* is a loss function that can measure the quality of function *f*, *R* is a regularization term defined over *f*, and *λ* is a regularization weight that captures the tradeoff between loss minimization and regularization (Bishop, 2006). In training the model, 75% of the observations in the statistical sample (*N* = 4,840) were randomly chosen to form the training set (*N* = 3,630), and the remaining 25% of the sample (*n* = 1,210) was used as the (hold-out) test set. All analysis was performed in R (version 3.6.1).

## 4. Results

### 4.1 Understanding the variations in alcohol drinking behavior explained by conventional and genetic factors

We first examined whether genetic factors are associated with participants’ *general* drinking behavior (as opposed to their drinking behavior related to a specific alcoholic product), which is measured by a binary variable indicating one’s drinking-or-not status in general (where 0 and 1 represent *current non-drinkers* and *current drinkers*, respectively). Fig. 2 compares the variances of participants’ alcohol use status explained by different sets of factors in the full sample (*left*), males subsample (*centre*) and females subsample (*right*). For the full sample, individual-level socio-demographic characteristics and genetic factors explained, respectively, 21.5% and 16.0% (of which single genetic factors explained 14.3% and PGSs explained 1.7%) of the total variance in alcohol use in general. Nevertheless, for males, genetic factors explained a significantly larger amount of variance (23.8%) compared to conventional socio-demographic factors (9.2%). Interestingly, an opposite pattern was observed for females: socio-demographic factors (21.6%) explained more variance than genetic factors (14.0%), and within the realm of genetic factors, polygenic scores (9.8%) outperformed single genetic markers (4.2%). This is consistent with previous findings from behavioral genetics that the genetic effects for alcohol use were more potent in males than in females using twinning and adoption designs (McGue et al., 1992; Verhulst et al., 2015). Two important implications can be derived from these findings. First, both conventional socio-demographic characteristics and novel genetic factors at the individual level are crucial for one’s alcohol drinking decisions. Second, and more importantly, in making these decisions, the combined effect of genetic factors sometimes outweighs that of socio-demographic characteristics, although substantial differences existed across gender groups.

**Fig. 2.**
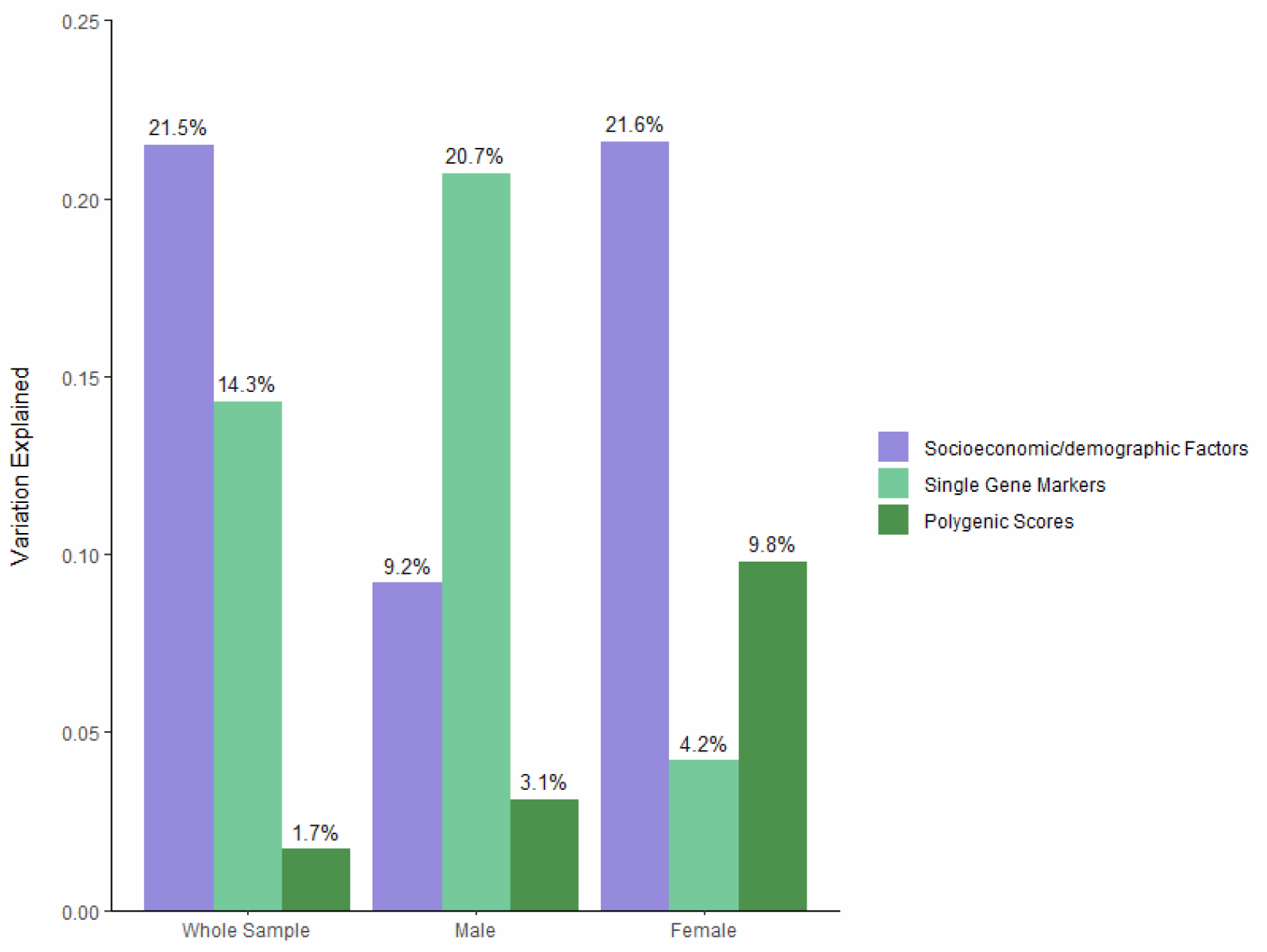
Variance explained by genetic and non-genetic factors. Source: Author’s survey.

### 4.2 Relationships between choice of alcohol and conventional/genetic factors

We then explored how conventional socioeconomic/demographic (i.e., non-genetic) factors and novel genetic factors are correlated with participants’ *particular* choice among four alcoholic alternatives (i.e., alcopop, beer, wine, and spirit, with an opt-out option) based on a DCE. Fig. 3 (a) and (b) present statistically significant correlation coefficients (*P* < 0.05) between alcohol alternatives with non-genetic and genetic factors, respectively. For example, male is positively correlated with choosing beer (*ρ* = 0.1038, *P* < 0.0001) and spirit (*ρ* = 0.1288, *P* < 0.0001), whereas age is negatively correlated with choosing beer (*ρ* = −0.0584, *P* = 0.0002) and wine (*ρ* = −0.0576, *P* = 0.0002); among genetic factors, *ALDH2* rs671 is negatively correlated with choosing beer (*ρ* = −0.0871, *P* < 0.0001) and spirit (*ρ* = −0.1157, *P* < 0.0001), while *OXTR* rs53576 is positively correlated with choosing alcopop (*ρ* = 0.0793, *P* < 0.0001) and beer (*ρ* = 0.0371, *P* = 0.0162).

**Fig. 3.**
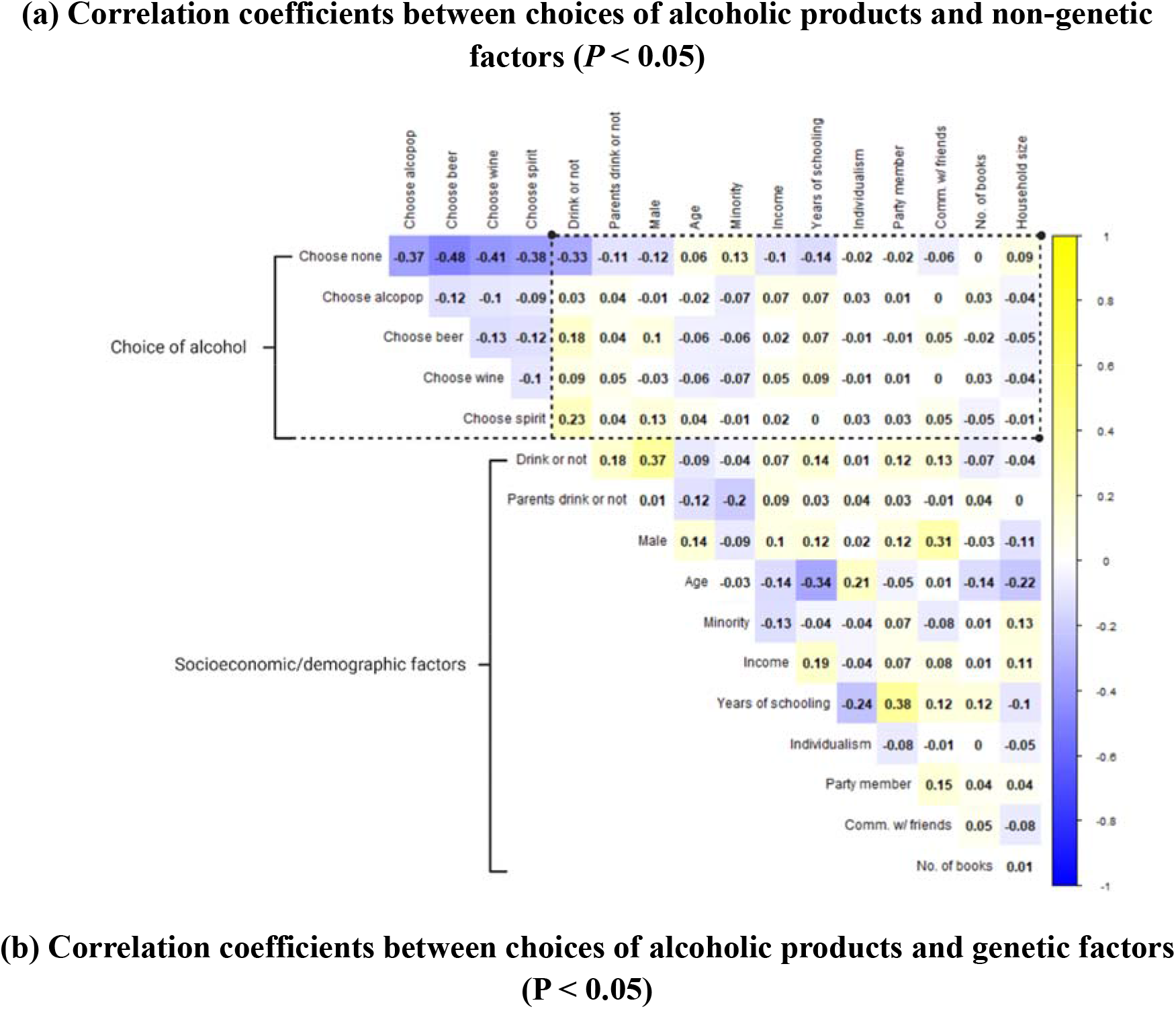
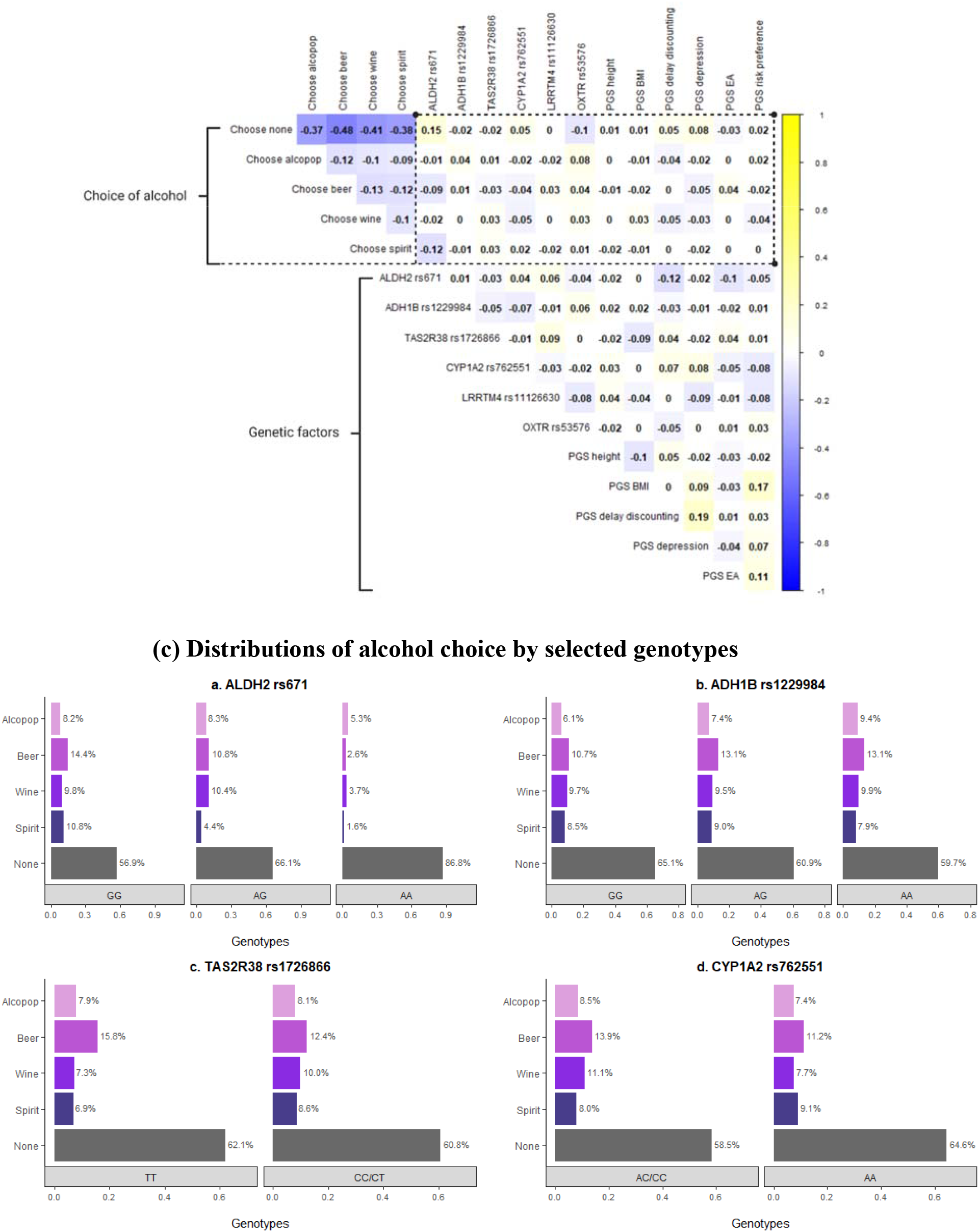
Relationships between alcoholic product choice and genetic/non-genetic factors. (a) The heat map depicts the correlation coefficients between choices of alcoholic products with each socioeconomic/demographic factor. All statistically insignificant correlation coefficients (*P* ≥ 0.05) were set to 0. (b) The heat map depicts the correlation coefficients between choices of alcoholic products with each genetic factor. All statistically insignificant correlation coefficients (*P* ≥ 0.05) were set to 0. (c) The distribution of alcohol choice by selected genetic variants. Source: Author’s survey and discrete choice experiment.

Fig. 3 (c) further demonstrates participants’ choice frequencies of alcoholic products (including the opt-out option) by genotypes of *ALDH2* rs671 (panel *a*), *ADH1B* rs1229984 (panel *b), TAS2R38* rs1726866 (panel *c*), and *CYP1A2* rs762551 (panel *d*), providing additional evidence that individuals’ choice of alcoholic products varies across different genotypes. Panel *a*, for example, suggests that participants are more likely to choose the opt-out option (in grey) as the number of effect alleles (A-allele) increases from 0 (genotype of GG, 56.9%) to 2 (genotype of AA, 86.8%). This is not surprising because the presence of A-allele in *ALDH2* rs671 has been found to significantly decrease the detoxification of acetaldehyde generated during alcohol metabolism in humans, resulting in the accumulation of acetaldehyde and several alcohol-intolerance/aversive reactions, such as alcohol flush, nausea, and rapid heartbeat (Peng et al., 2019; Edenberg and McClintick, 2018). Furthermore, for those participants who carry two effect alleles (genotype of AA), only 1.6% of them voluntarily chose spirits that have the highest ethanol concentration, considerably lower than among participants who do not carry any effect allele (genotype of GG, 10.8%) or carry one effect allele (genotype of AG, 4.4%). Overall, the variability in preference and choice to alcoholic beverages across participants can be at least partially explained by variations in their genetic endowments, lending initial support to the hypothesis that genetic factors contribute to individuals’ food choice decisions.

### 4.3 Model performance without and with genetic factors

To evaluate the relevance and strength of genetic factors in predicting individual choice of alcoholic products compared with conventional predictors, we developed five models. Model 1 (the Benchmark Model) only included product attributes as the base predictors. Models 2, 3 and 4 sequentially added socio-demographic features, single genetic markers, and polygenic scores, respectively, as additional predictors. Model 5 further included interaction terms of genetic and non-genetic characteristics as additional Gene-by-Environment (G-by-E) features. By comparing the performances of Model 1/2 and Model 3/4/5, we can empirically assess the contribution of genetic features versus that of conventional product/socio-demographic features to the prediction accuracy of individual alcohol choice.

After initial comparison of various machine learning algorithms (i.e., neural network, random forest, k-nearest neighbors, support vector machine, gradient boosting machine, extreme gradient boosting), the best performing classifier was found to be the extreme gradient boosting (XGBoost) classification model (Chen and Guestrin, 2016), which was thus chosen as our prediction model and testing ground. We trained the XGBoost model on the training sample, comprised of 75% observations randomly selected from the original sample, and tested the model on the remaining 25% of observations to maximize the generalizability of the model. To avoid overfitting, we implemented 10-times repeated tenfold cross-validation (CV) on the training dataset. Model performance was then evaluated based on Receiver Operator Characteristic (ROC) curves (Fig. 4) of the test dataset. A ROC curve is a graphical presentation of the ability of a model to distinguish among individuals with different alcohol choices, and is more informative than prediction accuracy for imbalanced choice data (Mandrekar, 2010). The area under the ROC curve (AUC) can be interpreted as the probability that the model will correctly assign to a randomly chosen participant, which ranges from 0.5 (worst classifier) to 1.0 (perfect classifier).

**Fig. 4.**
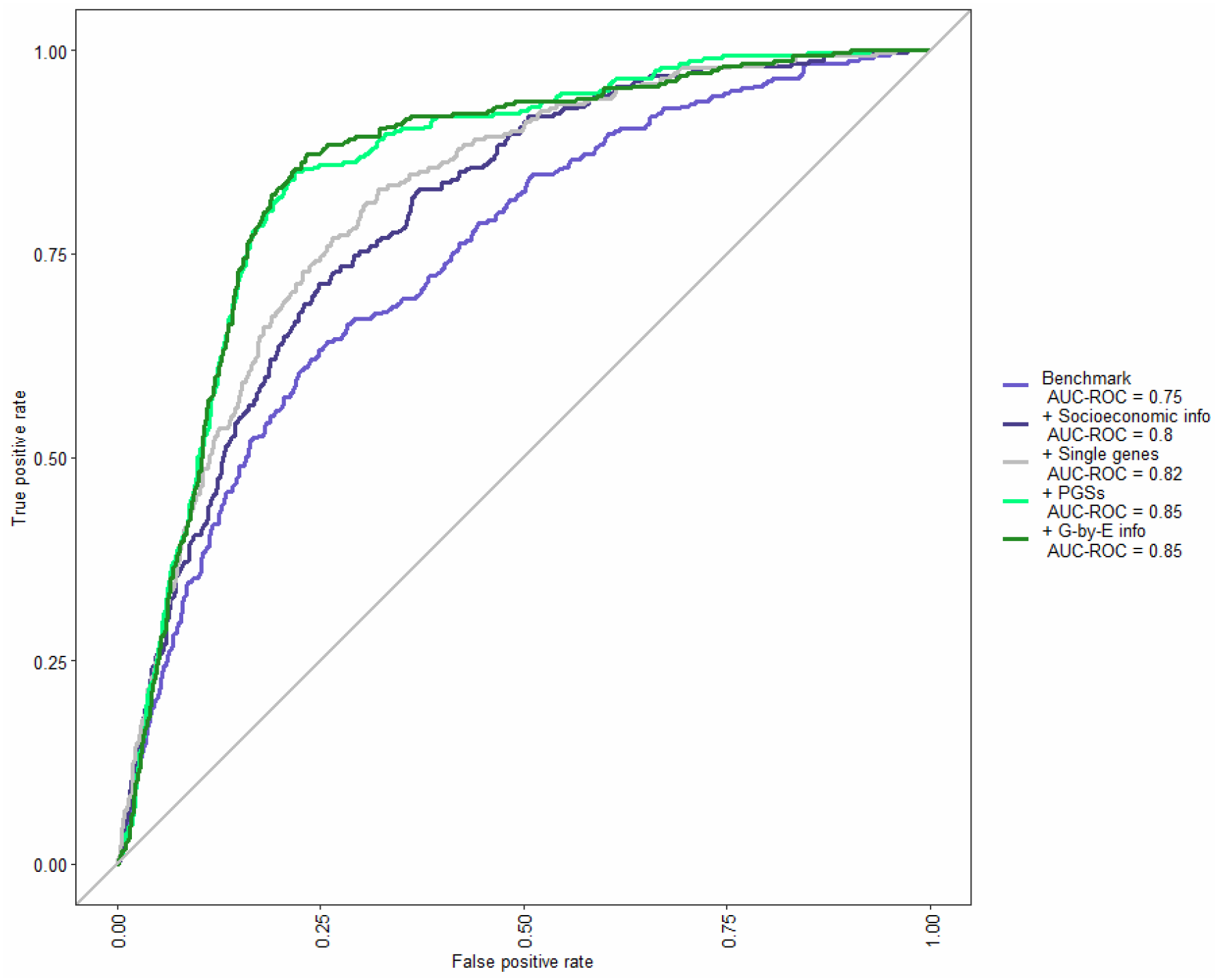
ROC curves for XGBoost classifiers with different sets of features. Source: Author’s analysis.

Fig. 4 displays ROC curves of all five models as mentioned earlier. The Benchmark XGBoost Model that includes only product attributes achieved an AUC-ROC of 0.75 and an accuracy of 63.4%, which are 19% and 17% higher than that of a traditional multinomial logit model (MNL) that uses the same set of explanatory variables (AUC-ROC_MNL_ = 0.63, AccuracyMNL = 54.1%).^1^ This confirms that machine learning-based classification models can significantly improve the efficiency and accuracy of individual choice prediction, compared with traditional discrete-choice model-based approaches. Three findings emerge when sequentially adding socioeconomic/demographic features (Model 2), single gene markers (Model 3), polygenic scores (Model 4), and G-by-E interactions (Model 5) as additional predictors. First, as expected, the predicting ability of Model 2 with both product attributes and individual-level socioeconomic/demographic features (AUC-ROC_Model2_ = 0.8, Accuracy_Model2_ = 69.6%) exceeds that of Model 1 with product features alone. Second, Model 4 performs the best in which both single genetic variants and PGSs are parts of the feature set, with an AUC-ROC reaching 0.85. Third, we observed no significant performance improvement when further adding G-by-E interactions in Model 5. Therefore, to minimize the risk of overfitting due to the inclusion of too many less useful features (Reunanen, 2003), the best performing classifier of Model 4 was selected as the final model, and the prediction accuracy of individual choice achieved 74.7% (95% CI: 73.1%-76.2%) from the test dataset.

### 4.4 Comparing the relative importance of features

To further explore the relative significance of each feature using the machine learning-based classifier, we conducted a feature importance analysis. The results are presented in Fig. 5. All relative importance scores are normalized by the score of the price (set to be 100) to show each feature’s relative contribution to the prediction of an individual’s choice of alcoholic beverages. For ease of presentation, the three subsets of individual-level features are presented in different colors: (A) socioeconomic/demographic factors (18 features), shown in purple; (B) single genetic variants (six features), shown in light green; and (C) polygenic scores (seven features), shown in dark green.

**Fig. 5.**
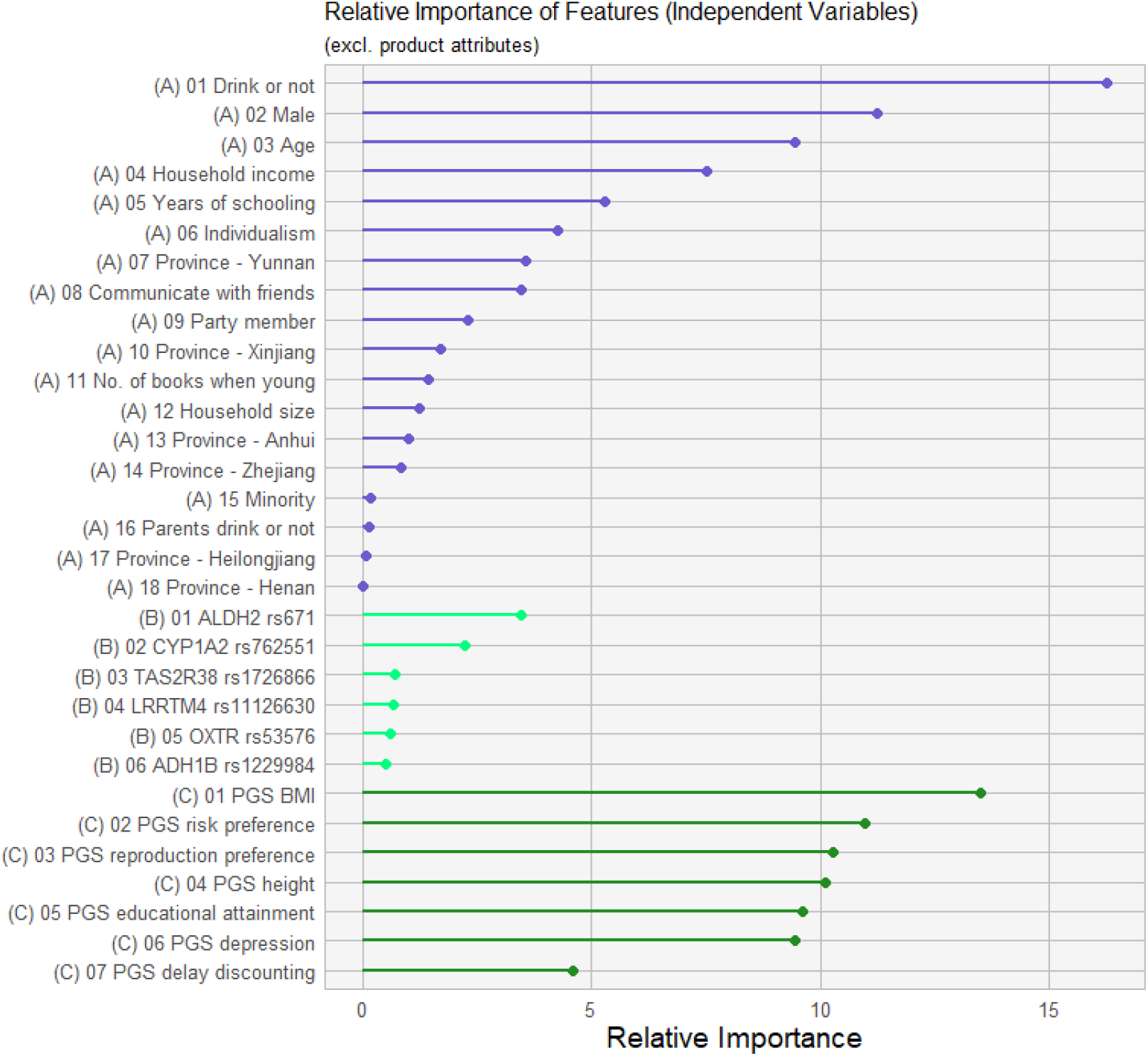
Feature importance results obtained from XGBoost. Source: Author’ calculation.

Among the 18 conventional socio-demographic features, the general behavior of drinking or not contributes the most to the classification, followed by gender, age, and household income. Among the six single genetic features, *ALDH2* rs671 and *CYP1A2* rs762551 are the top contributing features, which are more important than some of the socio-demographic factors, such as party membership and ethnic minority. Among the seven features of polygenic scores, the PGS of BMI is the most critical in predicting individual alcohol choice, which surprisingly outperforms most of the socio-demographic features (except drinking or not in general), including gender, age, income, and educational attainment that are known as important factors for explaining consumer heterogeneity in food preference and choice (Shepherd, 2005; Kearney et al., 2000). Simply put, not only does the feature importance analysis support the association between genetic factors and food choice decisions at the individual level, but it also demonstrates that the predictive strength of genetic factors is comparable to several well-documented socio-demographic characteristics.

## 5. Conclusions

The complex and heterogeneous nature of human behavior has raised many challenges in related disciplines. Our analysis demonstrates that genetic factors are not only relevant to individuals’ food choice decisions, but also play a similarly critical role in the prediction of individual choice of alcoholic beverages compared to the conventional socio-demographic characteristics. By creating the best well-performing machine learning-based classification model with both socio-demographic and genetic features, we achieved the highest accuracy of 74.7% and AUC-ROC of 0.85, which can be considered as an excellent classifier in practice (Mandrekar, 2010). Incorporating genetic factors to the MLC predictive model improves both of the accuracy (by 5.1 percentage points) and the AUC-ROC (by 0.05). Furthermore, among all genetic factors, the polygenic score of BMI has the highest predictive power for individual alcohol choice, comparable to the contribution of several well-known socio-demographic characteristics, such as gender, age and income. Overall, not only are our findings in accordance with the small but emerging evidence that genetic factors can influence the general food intake/dietary composition of an individual, but they also demonstrate the contribution of genetic features to individual preference/choice among similar alternatives that is subtler, warranting further studies of consumer behavior with the integration of individual genotyping information. A food choice model adapted from Connors et al. (2001), presented in Fig. 6, illustrates how personal factors (i.e., demographic, socioeconomic, and genetic factors) would affect the food choice decision-making process through an individual’s unique personal food system (Feeney et al., 2011). As recent evidence has shown that general dietary interventions rarely fit the population as a whole, researchers may benefit from integrating genetic profiles to develop more effective interventions that are tailored to the personalized food demand of targeted individuals (Daviet et al., 2020; Garcia-Perez et al., 2020; Zeevi et al., 2015; Clarke et al., 2015; Shepherd, 2005).

**Fig. 6.**
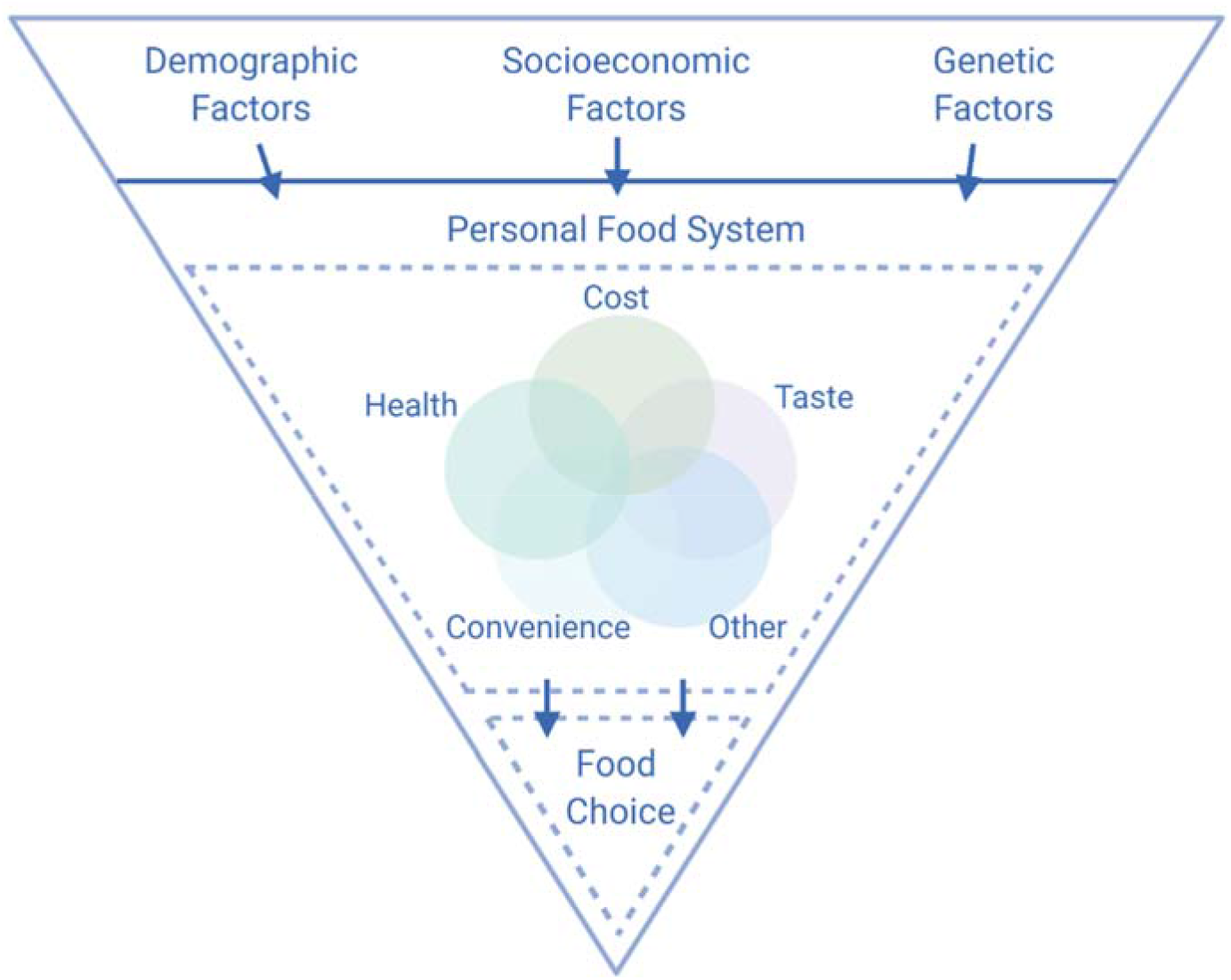
How personal factors influence an individual’s unique food system and choice decisions. Source: Adapted from Connors et al. (2001) and Feeney et al. (2011).

Before closing, we note several limitations in the current study. First, the HCG sample used in this study may not be representative of the entire Chinese population. More work would be required to confirm the external validity of our findings in other populations. Second, the current study was based on a sample of only 484 participants, which may lack statistical power to detect the contribution of other genetic factors to one’s food choice, due to its relatively small sample size. Third, our DCE was focused on alcoholic products; more work would be needed to generalize our findings to other food categories. As a future step, further integration of individual genotyping data into studies of consumer behavior and decision-making process, as well as a larger and more population-representative samples, would provide a better understanding of the delicate but nontrivial effects of genetic predisposition on the individual decision-making process.

The current study provides two distinctive contributions. First, by integrating individual-level genotyping data with the advanced economic research design of discrete choice experiments, we directly show the importance of genetic factors for their contributions to individual choice decisions of differentiated food products for the first time in the literature. Second, by employing machine learning-based classification models, we systematically and empirically proved that genetic factors could substantially improve the prediction of individual food choice of alcoholic beverages. We unpack the relative importance of genetic features that are comparable to several well-documented socio-demographic characteristics. Overall, this study broadens our understanding of how individuals make food choice decisions and illustrates a more effective framework for predicting human decisions/behaviors at the individual level. As more large-scale socioeconomic surveys around the world started to collect and disclose genotyping data (e.g., the Health and Retirement Study, the National Longitudinal Study of Adolescent to Adult Health), the integration of genetic features into economic studies has become more and more cost-effective, and we hope that our findings can inspire more work in decision-making domains of interest to social scientists.

## Footnotes

1 The classification accuracy is defined as the ratio of correct predictions to total predictions made.

